# A Truly Injectable Neural Stimulation Electrode Made from an In-Body Curing Polymer/Metal Composite

**DOI:** 10.1101/584995

**Authors:** James K. Trevathan, Ian W. Baumgart, Evan N. Nicolai, Brian A. Gosink, Anders J. Asp, Megan L. Settell, Shyam R. Polaconda, Kevin D. Malerick, Sarah K. Brodnick, Weifeng Zeng, Bruce E. Knudsen, Andrea L. McConico, Zachary Sanger, Jannifer H. Lee, Johnathon M. Aho, Aaron J. Suminski, Erika K. Ross, J. Luis Lujan, Douglas J. Weber, Justin C. Williams, Manfred Franke, Kip A. Ludwig, Andrew J. Shoffstall

**Affiliations:** Department of Biomedical Engineering, University of Wisconsin-Madison, Madison, WI, USA; Mayo Clinic Graduate School of Biomedical Sciences, Mayo Clinic, Rochester, MN, USA; Department of Neurosurgery, University of Wisconsin-Madison, Madison, WI, USA; Department of Biomedical Engineering, Case Western Reserve University, Cleveland, OH, USA; Department of Neurologic Surgery, Mayo Clinic, Rochester, MN, USA; Division of General Thoracic Surgery, Mayo Clinic, Rochester, MN, USA; Physiology and Biomedical Engineering, Mayo Clinic, Rochester, MN, USA; Department of Bioengineering, University of Pittsburgh, Pittsburgh, PA, USA; Neuronoff Inc., Valencia, CA, USA; Advanced Platform Technologies Center, Louis Stokes Cleveland Veterans Affairs Medical Center, Cleveland, OH, USA

**Keywords:** Neuromodulation, Peripheral Nerve, Neural Interface, Injectrode, Biomaterial

## Abstract

Implanted neural stimulation and recording devices hold vast potential to treat a variety of neurological conditions, but the invasiveness, complexity, and cost of the implantation procedure greatly reduce access to an otherwise promising therapeutic approach. To address this need, we have developed a novel electrode that begins as an uncured, flowable pre-polymer that can be injected around a neuroanatomical target to minimize surgical manipulation. Referred to as the Injectrode, the electrode conforms to target structures forming an electrically conductive interface which is orders of magnitude less stiff than conventional neuromodulation electrodes. To validate the Injectrode, we performed detailed electrochemical and microscopy characterization of its material properties and validated the feasibility of using it to electrically stimulate the nervous system in rats and swine. The silicone-metal-particle composite performed very similarly to pure wire of the same metal (silver) in all measures, including exhibiting a favorable cathodic charge storage capacity (CSC_C_) and charge injection limits compared to the clinical LivaNova stimulation electrode and silver wire electrodes. By virtue of being simpler than traditional electrode designs, less invasive, and more cost-effective, the Injectrode has the potential to increase the adoption of neuromodulation therapies for existing and new indications.

## 1. Introduction

Electrical stimulation of the peripheral nervous system, often known as neuromodulation, Bioelectronic Medicines^[1]^ or Electroceuticals^[2, 3]^, is an increasingly prevalent clinical therapy. In recent years, the Federal Food and Drug Administration (FDA) has approved market applications for myriad of new indications for Bioelectronic Medicines including diverse conditions such as obesity, sleep apnea, migraine, opioid withdrawal symptoms, dry eye, essential tremor and hypertension. These supplement traditional market indications for Bioelectronic Medicines such as pain and overactive bladder.^[4, 5]^ Additional areas of research currently beginning early stage clinical testing include the use of therapeutic electrical stimulation to treat inflammation, metabolism, and endocrine disorders.^[6–9]^ The increasing prevalence of Bioelectronic Medicines has led to a renewed interest in both developing novel devices as well as improving the mechanistic understanding of how neural interface technologies interact with the nervous system. This has resulted in numerous large-scale government funding programs such as the White House BRAIN Initiative, the NIH SPARC Program, and the DARPA HAPTIX and ElectRx Programs.^[10]^

Existing electrical stimulation devices designed for integrating with the human nervous system can be separated into non-invasive devices, minimally invasive hybrid strategies, and invasive devices. Non-invasive devices apply an electrical stimulation waveform through electrodes placed on the surface of the skin, which are typically intended to electrically interface with deeper neural structures (Figure 1B). Although some impressive therapeutic responses have been demonstrated using non-invasive stimulation paradigms targeted to specific nerves, the unintended stimulation of nerve and muscle fibers superficial to the deep target often lead to therapy limiting side-effects.^[11–13]^ Additionally, the electric field falls off rapidly with distance from the stimulating electrode, which limits the depth at which structures can be reliably stimulated, at least without intolerable activation of cutaneous pain fibers.^[14]^

**Figure 1.**
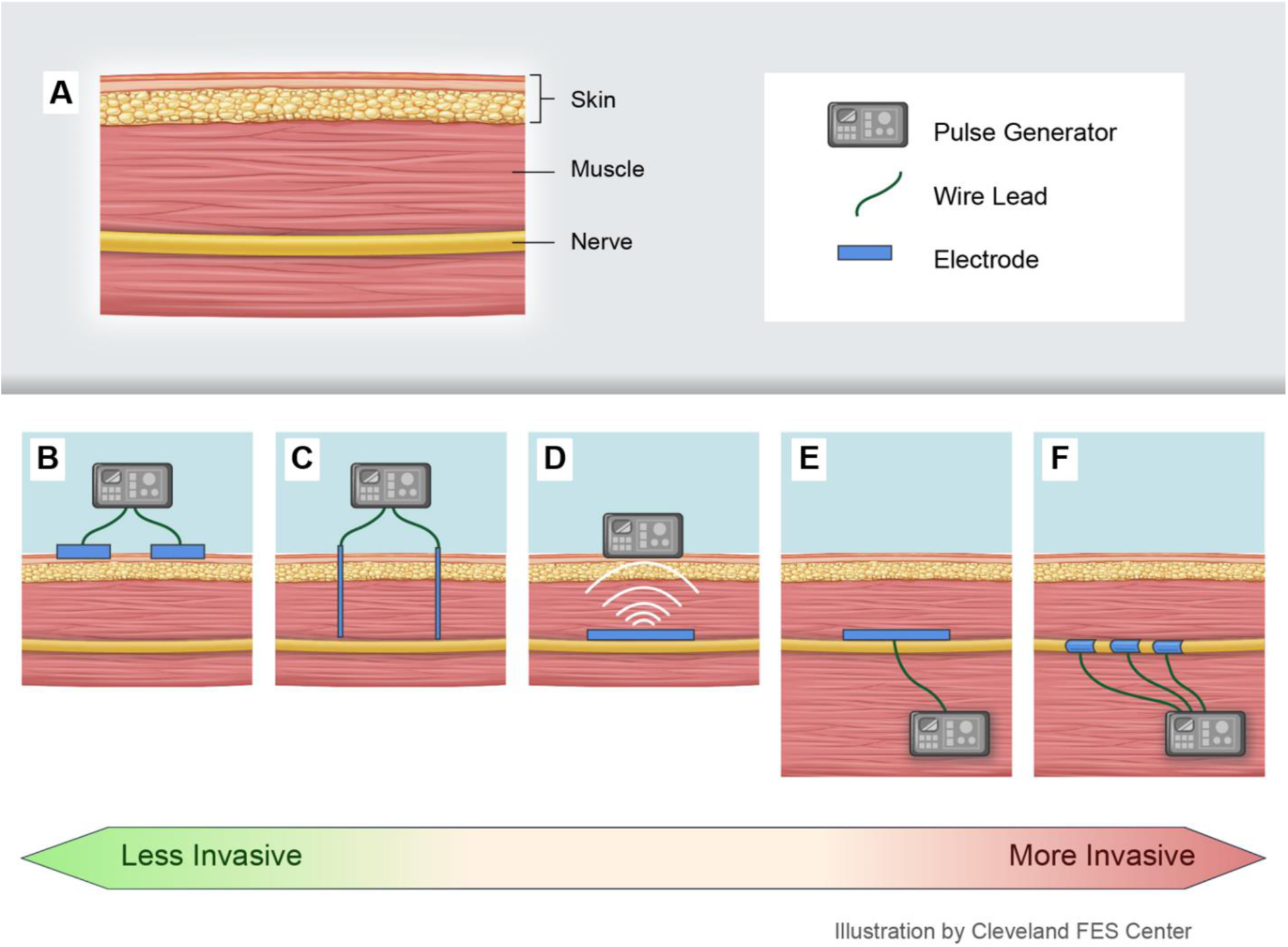
Peripheral nerve stimulation electrode types from least to most invasive, left-to-right. **A.** simplified tissue schematic showing distinct layers of the skin muscle and nerve. **B.** transcutaneous electric nerve stimulation (TENS), **C.** percutaneous electric nerve stimulation (PENS), **D.** injected wireless stimulator (e.g., BION® device), **E**. implanted nerve stimulator (e.g. spinal cord paddle electrode), **F.** implanted cuff electrodes (e.g. vagus nerve stimulator).

Given the limitations of non-invasive stimulation strategies, implantable neural stimulation electrodes are commonly used to provide more precise activation of a nerve trunk, ganglia, nuclei or other targets of interest. One of the most common implantable interfaces to stimulate the peripheral nervous system is the nerve cuff, which consists of one or multiple electrodes positioned on an insulating backer that is surgically placed with the electrode contact(s) surrounding the epineural surface of a nerve trunk (Figure 1F).^[15]^ Although this approach can provide very specific target engagement, the placement of nerve cuffs requires surgical dissection down to the nerve trunk of interest through skin, muscle, smaller nerves and microvasculature. Moreover, cuff implants typically requires 360-degree dissection around the nerve trunk of interest to facilitate placement around the nerve. Additionally, stimulation leads must be tunneled to an implantable pulse generator (IPG) that has a limited battery life. Multiple points of failure and the complexity of the implantation procedure increases both surgical risk and costs, and may contribute to a large amount of variability in the therapeutic effectiveness of peripheral neuromodulation devices. For example, a 152 patient clinical study recently compared invasive Dorsal Root Ganglion (DRG) stimulation to traditional epidural spinal cord stimulation (SCS) for intractable pain. 81.2% of the DRG patients received a greater than 50% decrease in back pain vs 55.7% with SCS, a significant difference. However, the adverse event rate related to the neural stimulator/device or the implant procedure was significantly higher in the DRG arm (36.8% to 26.4% and 46.1% to 26.3%, respectively). As both DRG and SCS require a complex electrode/lead/IPG system, implant costs ranging from $32k to $58k and annual maintenance costs ranging from $5k to $20k also pose significant barriers to treatment.^[16]^

Practically speaking, stimulation device selection involves carefully balancing trade-offs between efficacy and safety – including implantation invasiveness, efficacy, robustness and the cost of the procedure. Although stimulation and implantation strategies spanning the spectrum from non-invasive to fully implanted (Figure 1) are available, there remains an unmet need for neural interface technologies that can be implanted through a minimally invasive procedure but can maintain a robust connection with complex and difficult to surgically dissect peripheral neuroanatomy.

To meet this need, we have developed a novel electrode, which we call the ‘Injectrode’ (Figure 2). The Injectrode is flowable as a pre-polymer and is injected via a syringe where it cures to form a highly conforming, compliant neural electrode *in vivo.* By flowing around the target neuroanatomy, the Injectrode can conform to a variety of targets to form different neural interfaces. For instance, the neural engagement of an invasive cuff electrode could be mimicked by injection of the electrode into the sheath around a nerve trunk. Alternatively, the Injectrode can be used to stimulate complex neural structures such a nerve plexus or those inside a foramen that may be very difficult to target with traditional cuff electrodes. Through the injection of this electrode, neural interfaces can be created using a minimally invasive surgical approach that avoids the risks of surgical complication associated with open cut-downs and extensive dissection. Although, the Injectrode is eventually intended to be delivered in a minimally invasive fashion in a clinical population, the goal of this manuscript was to characterize the electrochemical properties of the composite material to demonstrate potential as a neural stimulation electrode as well as to show proof of concept stimulation of complex neuroanatomy in small and large animal models. Development of methods for precise and reproducible administration of the Injectrode to form functional multi-contact neural interfaces in a clinical setting are reserved for future work.

**Figure 2:**
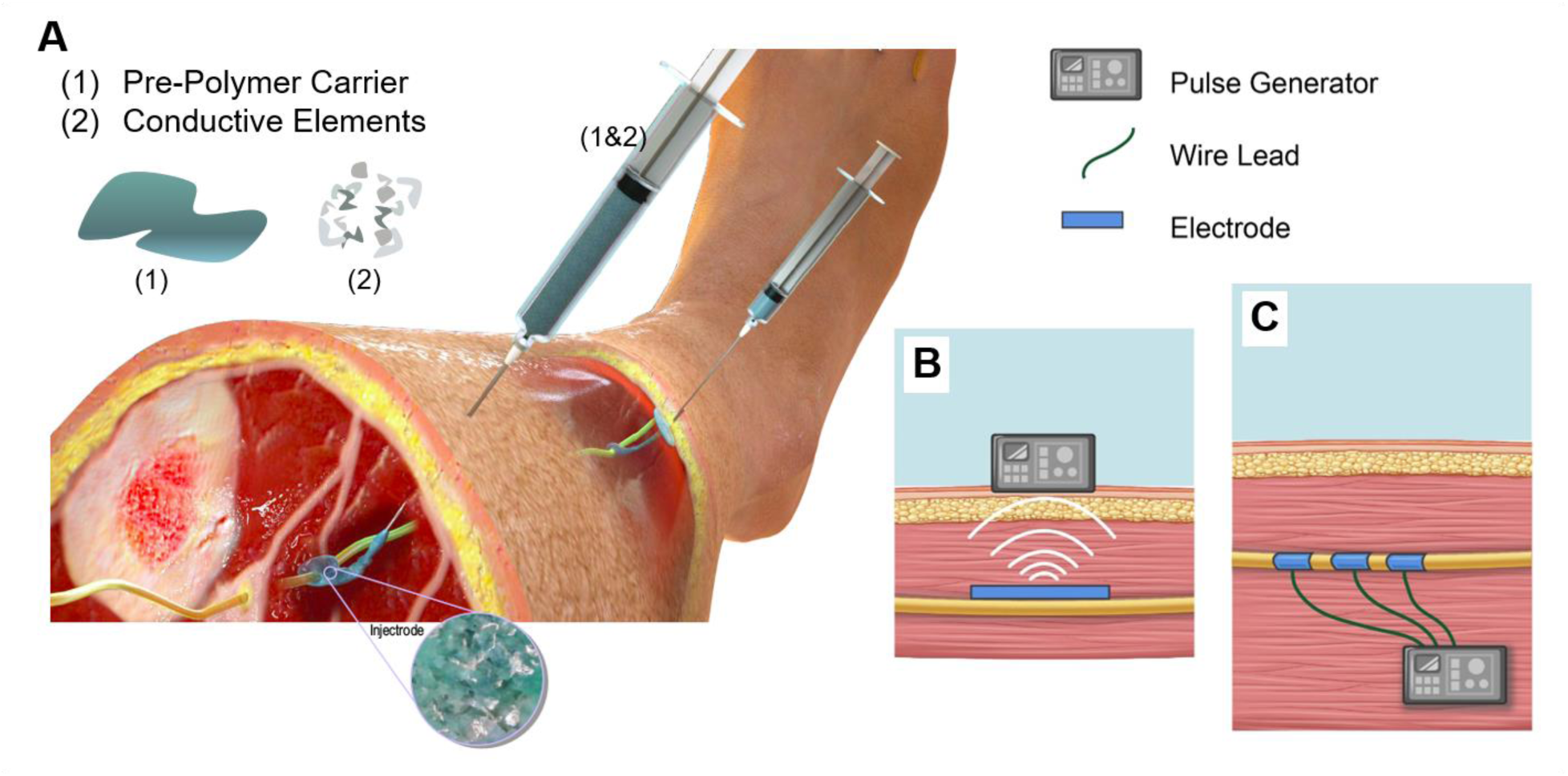
The Injectrode™ Concept. **A.** A composite material consisting of (1) a pre-polymer carrier and (2) conductive filler elements is injected into the body near a target nerve. The mixture cures in place to form an electrically conductive interface. **B.** wireless configuration (e.g., if connected with a RF antenna circuit), **C.** wired configuration (e.g. if directly connected to an IPG).

Herein, we present detailed electrochemical and microscopy characterization of material properties of the Injectrode. Additionally, we validate the feasibility of electrically stimulating peripheral nerves with an *in vivo* curing composite material in pre-clinical animal models. Visual assessments were made of the surface morphology of electrodes via light microscopy and scanning electron microscopy. Cyclic voltammetry, voltage excursions, and electrochemical impedance spectroscopy were used to directly compare the electrochemical properties of the Injectrode to those of pure silver wire, as well as clinical LivaNova Platinum/Iridium electrodes. Lastly, we also present proof of concept experiments in small and large animal models in order to demonstrate the feasibility of this material as an *in vivo* neural interface.

## 2. Materials and Methods

### 2.1. Materials and Equipment

All general supplies, chemicals, reagents and buffers were sourced from Sigma (St. Louis, MO) and used as received. Proprietary mixtures of two-part Pt-curing silicone elastomer and metallic silver flakes were prepared according to instructions from Neuronoff, Inc (Valencia, CA). Blend percentages are defined according to a weight/volume (w/v) nomenclature with respect to silver/silicone, where 80% w/v corresponds to 800 mg sliver flakes and 200 mg mixed silicone pre-polymer. The silver flakes (25-50 µm aggregate size) were purchased from Inframat Advanced Materials and used as received. The flakes were additionally pre-treated with (3-Glycidyloxypropyl)trimethoxysilane (GLYMO) to improve their incorporation into the silicone matrix according to manufacturer recommendations.

To control geometry for the benchtop characterization experiments, materials were mixed thoroughly and immediately transferred to cure in either a 96-well plate or a 1/16” inner diameter silicone tubing for incorporation into a working electrode, as shown in Figure 3. For the *in vivo* experiments, materials were mixed and immediately transferred to a syringe for injection (extrusion) around the nerve of interest.

**Figure 3:**
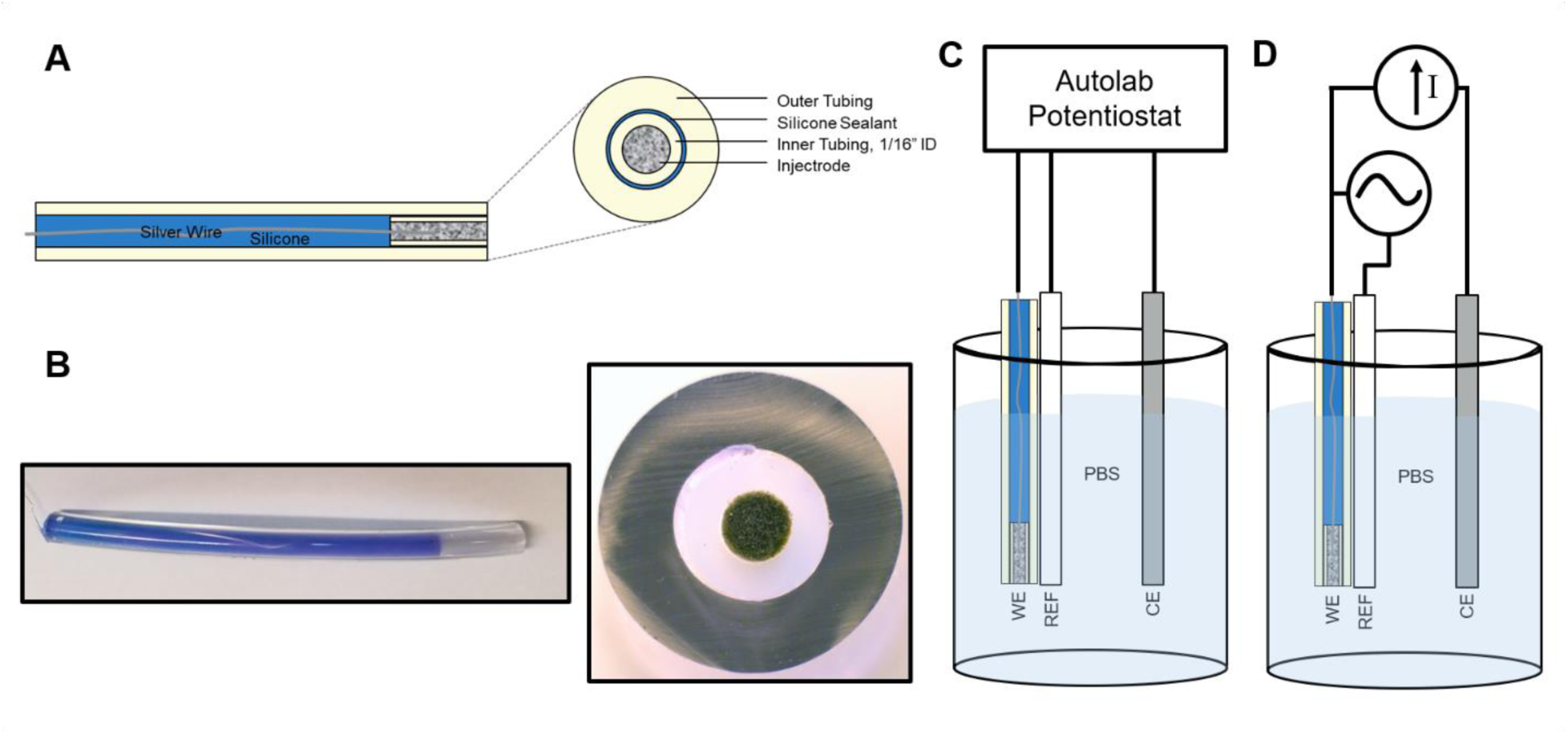
Working electrode fabrication schematic. **A.** A short segment of Injectrode material was filled into a 1/16” ID tubing and an uninsulated silver wire was inserted into one end. A larger ID tubing was placed over the smaller tubing to form an insulative outer sheath and filled with silicone. The constructed working electrode provides a consistent 1/16” diameter disk of Injectrode exposed at one end with a silver wire exposed at the other to provide a connection for performing electrochemical experiments **B.** (left) An image of a completed Injectrode sample fabricated into a working electrode. (right) Cross-section of working electrode tip showing the Injectrode sample with a controlled geometric surface area. **C.** Diagram of a three electrode electrochemical setup used to perform electrochemical testing. **D.** Diagram of electrochemical setup used for acquiring voltage transient data, with a current source delivering the waveform between the working electrode (WE) and counter electrode (CE) as well as an oscilloscope for measuring applied voltage with respect to a silver/silver chloride (Ag/AgCl) reference electrode (REF).

### 2.2. Electrochemical Characterization

Benchtop electrical characterization generally followed the techniques described in Stuart Cogan’s 2008 review of neural stimulation and recording electrodes.^[17–24]^ To achieve consistent electrochemical testing, working electrodes with tightly controlled geometric surface area were fabricated. A pre-polymer that was 80% (w/v) silver particles was mixed as previously described and immediately injected into a 1/16” inner diameter (ID) tube (labeled Inner Tubing in Figure 3), which was then cut into 2 cm segments. While curing, a 0.25 mm diameter silver wire was embedded approximately 1 cm deep into each Injectrode sample. After curing, the inner tubing piece was placed into a second piece of tubing (labeled Outer Tubing in Figure 3) chosen with an ID slightly larger than the outer diameter (OD) of the inner tubing. The electrode was then back-filled with silicone, which served to provide mechanical stability and insulate the silver wire. This yielded an electrode with a 1/16” diameter circular disc Injectrode material exposed at one end, which was used as a working electrode for electrochemical characterization experiments measuring cyclic voltammetry, electrochemical impedance spectroscopy, and voltage transients. A schematic of the fabricated working electrode is shown in Figure 3A and photographs showing both a working electrode sample and magnified cross-section of the electrode surface are presented in figure 3B.

For comparison of Injectrode samples to a pure silver electrode, silver working electrodes were fabricated by removing, under a microscope, insulation from 0.01” silver wire (A-M Systems, Carlsborg, WA) such that the total exposed surface area matched the geometric surface area of the Injectrode, which was 0.0197 cm^2^. In order to compare properties of the Injectrode to a clinically relevant electrode, a LivaNova PerenniaFLEX^®^ stimulation electrode was used.

Cyclic voltammetry and electrochemical impedance spectroscopy were measured in 0.01 M phosphate buffered saline (PBS) (NaCl 0.138 M; KCl - 0.0027 M; pH 7.4; Sigma, St. Louis, MO) using a three electrode cell consisting of a previously described Injectrode working electrode, a platinum sheet counter electrode (1 cm^2^ surface area; Metraohm, Herisau, Switzerland), and a single junction silver/silver chloride (Ag/AgCl) in 3M KCl reference electrode (BASi, West Lafayette, IN). Measurements were performed with an AutolabPGSTAT128N potentiostat (Metraohm).

Cyclic voltammetry was performed with a staircase sweep at 50 mV/s with a step size of 2.44 mV. 15 voltammograms were performed on each Injectrode sample and the final three voltammograms on each sample were averaged to obtain a single measurement. A new working electrode was used for each measurement to minimize effects from previous measurements on any given sample. In order to compare the electrochemical characteristics of the Injectrode to that of a silver electrode with the same surface area fabricated as previously described, extended voltammograms from −1.9 to 3.0V vs Ag/AgCl were also applied to both electrode types. In order to compare properties of the Injectrode to that of a standard platinum stimulating electrode a LivaNova PerenniaFLEX^®^ stimulation lead was used and cyclic voltammograms −0.62 to 0.9 V vs Ag/AgCl, a common range for platinum, were applied. Prior to the collection of voltammetry measurements, the LivaNova stimulation electrode was cleaned by sonicating for 30 minutes in isopropyl alcohol, then an additional 30 minutes in DI water, followed by anodic etching for 2 minutes at an applied potential of 2V vs. Ag/Cl in a 0.5M sulfuric acid solution. From the collected voltammograms the cathodal charge storage capacity (CSC_C_) was calculated for each electrode as the time integral of the cathodic current.

To characterize the impedance properties of the Injectrode and compare it to silver wire and platinum stimulation electrodes, electrochemical impedance spectroscopy (EIS) was performed with the same electrodes and electrochemical cell as was used for cyclic voltammetry. Electrochemical impedance spectroscopy was performed following the application of 20 voltammograms from −0.62 to 0.9V vs Ag/AgCl. Impedance spectroscopy was performed with 25mV sinusoidal waveforms applied with equal spacing on a logarithmic scale from 0.1 Hz to 100 kHz.

Lastly, voltage transient (VT) measurements were taken, once again, following the application of 20 voltammograms from −0.62 to 0.9V vs Ag/AgCl. A cathodic-leading charge-balanced biphasic waveform was used for the VT measurements with pulsewidths of 500 µs, and a 10 µs interpulse delay. Pulses were applied at 50 Hz and VT measurements were made following at least 1000 pulses in order to allow for stabilization. A Keithley 6221 current source (Tektronics, Beaverton, OR) was used for the application of pulse trains used for VTs. Using a Tektronics TBS1154 Oscilloscope (Tektronics, Beaverton, OR), VTs were recorded with the application of increasing amplitude pulses until the maximum cathodic polarization (E_mc_) was greater than 0.6V. To minimize the effects of noise, VT measurements were recorded as the average of 16 pulses using built in averaging on the oscilloscope.

### 2.3. Electrode Imaging

Light and electron microscopy of the electrode surface were performed on an unstimulated Injectrode sample from a working electrode that was fabricated as described in the previous section. In preparation for imaging, a cross-sectional slice (Figure 3B) of an Injectrode sample that had not undergone electrochemical testing or stimulation was cut with a scalpel.

Light microscopy was performed with an AmScope ZM-4TW3 stereo microscope, equipped with an 18 MP camera (AmScope, Irvine, CA). Images were converted to grayscale and contrast enhanced by histogram normalization in ImageJ (National Institutes of Health, Bethesda, MD). The slice was then platinum coated in preparation for SEM. SEM was performed using a Zeiss LEO 1530 (Zeiss International, Oberkochen, Germany) with an accelerating voltage of 3.0 kV.

### 2.4. Mechanical Testing

ASTM International has specified procedures for evaluating mechanical properties of thermoset plastics under Active Standard D412-16. Specified forms include flat specimens with uniform cross-sectional area, and dog bone shaped specimens. Molds were designed as closely as possible adhering to the guidelines for creating flat dog bone shaped specimens in SolidWorks. The molds were then cut from acrylic sheets using a laser cutter at ThinkBox (Case Western Reserve University). The mechanical properties of the cured electrodes were then tested using a pneumatically driven universal testing machine (Enduratec) located in the CWRU Department of Materials Science & Engineering Advanced Manufacturing and Mechanical Reliability Center (AMMRC). Specimens (n=4, independently mixed) were placed into the grips of the testing machine pulled under tension at a rate of 0.5 mm/s. Grip spacing was set to 2 cm. The first sample was tested to 140% strain and did not fail. Therefore, we increased the maximum strain for subsequent samples, which were pulled to a final extension of 6 cm (200% strain) or failure.

### 2.5. Acute Rat Brachial Plexus Stimulation

All animal experiments were approved by the Mayo Clinic Institutional Animal Care and Use Committee (Rochester, MN). Rats were anesthetized with isoflurane, using 5% isoflurane in pure oxygen for induction and 1-2% for maintenance over the duration of the experiment. The compound motor nerve branches of the rat brachial plexus were exposed and freed from surrounding tissues via careful dissection through the pectoralis muscles. Components of the Injectrode composite were combined in a syringe and mixed for 30 seconds to make the flowable pre-polymer, as previously described. The pre-polymer mixture was injected within 1-2 minutes, to create two circumferential ‘cuff’ electrodes in a bipolar configuration with 5 mm spacing, around the exposed rat brachial plexus. Silver wires were inserted into the composite during curing to make direct wired connections to the cuffs. For the purposes of this proof of concept experiment testing the Injectrode in an open surgical site, Kwik-Cast (World Precision Instruments, Sarasota, FL) was used to insulate between and around the two electrodes.

A biphasic charge balanced waveform at 30 Hz (200 µs pulse width, 5mA amplitude) was applied using Master-8 stimulator (A.M.P.I., Jerusalem, Isreal) with Iso-Flex stimulus isolators (A.M.P.I., Jerusalem, Isreal). Effect of stimulation was observed visually as large muscle contractions of the forelimb. Approximate joint angles were calculated using post-hoc video analysis. In brief, still images were captured from the video every 5 frames (30 fps), and analyzed in ImageJ. Two lines were drawn through the joint. The angle was measured using ImageJ’s built-in tools.

### 2.6. Acute Swine Vagus Nerve Stimulation

Pigs (weight 35-40 kg range) were anesthetized using Telazol/Xylazine for induction followed by maintenance with 1-2% isoflurane for the duration of the experiment. The vagus nerve was carefully dissected free from surrounding tissue and a commercially available bipolar cuff (LivaNova) was placed on the nerve approximately 0.5 to 1.0 cm inferior to the nodose ganglion and 1.0 to 1.5 cm from the carotid bifurication. The cuffs were placed on the ventromedial aspect of the vagus nerve.

A stimulation current-dose-titration curve was generated, with a biphasic charge balanced waveform at 30 Hz (200 µs cathodic pulse width), ranging from 0-5mA. Heart rate was monitored with a pulse oximeter and recorded.

To compare results obtained during stimulation with the Injectrode to those obtained using clinical LivaNova electrodes, cuff-like electrodes were fabricated using Injectrode material extruded onto a stainless-steel mesh with an insulated backing. The same stimulation current-dose-titration curve was generated and compared to the results obtained while stimulating with the LivaNova electrode.

## 3. Results and Discussion

Initial benchtop impedance measurements were undertaken to ascertain the percolation threshold of the material mixtures, which was found to occur at approximately 65% w/v mix with respect to silver content. (Figure S1). Based on these results, an 80% w/v with respect to silver content was selected to provide a wide margin of error and used for all of the electrochemical and in vivo testing presented in this manuscript.

### 3.1. Electrode Imaging

Light microscopy and electron microscopy were performed in order to assess the surface properties of the material. For both types of imaging a thin slice of material was cut from the surface of an Injectrode working electrode sample that had not undergone any electrochemical testing or stimulation. Imaging revealed that the silver particles were well distributed throughout the silicone matrix (Figure 4). Electron microscopy showed surface porosity (Figure 4C,D) as well as a complex arrangement of silver particles within in the silicone matrix.

**Figure 4:**
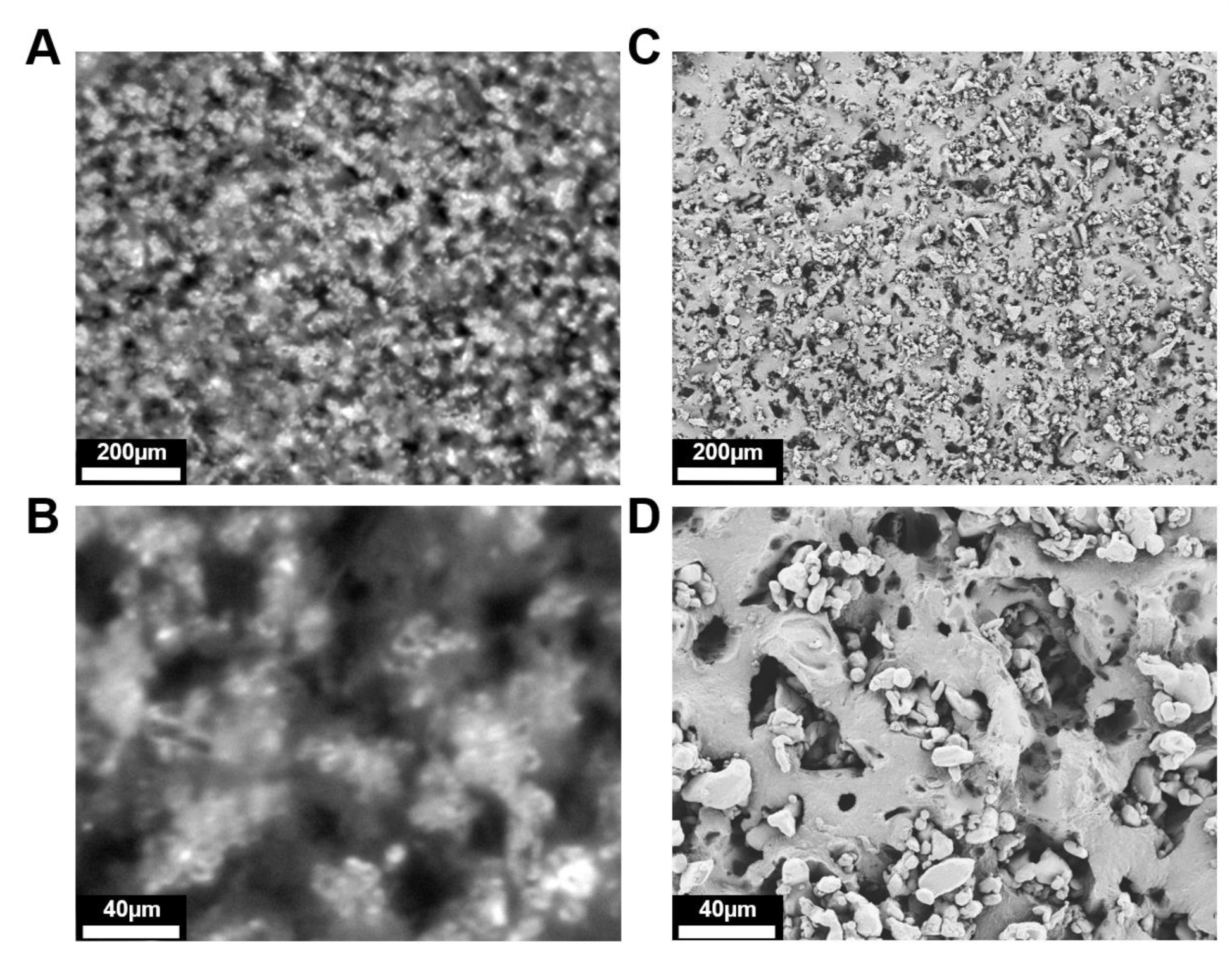
Light microscopy and SEM of the surface of an Injectrode. A-B. Light microscopy of the Injectrode surface showing distribution of silver particles throughout the sample. C-D. Scanning electron microscopy (SEM) at comparable magnification showing sample topography and revealing porosity of the polymer matrix.

The porosity of the silicone matrix may act to increase electrolyte permeability, which could enable electrode/electrolyte interactions with particles deeper within the silicone matrix. This in turn would increase the fractal dimension of the material and thus the effective electrochemical surface area. For these reasons, the porosity of the material may be important to the functional capabilities of the Injectrode as a stimulation electrode particularly with respect to increasing the effective surface area available for charge injection. Thus, modifying the porosity of the cured Injectrode is one factor that could potentially be controlled in order to improving the stimulation characteristics of the electrode.

### 3.2. Electrochemical Characterization: Cyclic Voltammetry

Cyclic voltammetry is a classical electrochemical method for the characterization of stimulation electrodes. It provides information regarding the electrode interface under electrical load, the electrochemical conversion of species within the solution, and on transient changes due to oxidation or other processes at the electrode surface.^[17]^

For assessment of Pt/Ir electrodes, performing slow scan (50 mV/s) cyclic voltammetry sweeps in PBS over a voltage range known as the water window; the window between the voltages at which evolution of hydrogen and oxygen occur is common practice.^[17, 25]^ Commonly used values for assessment of Pt and Pt/Ir electrodes include −0.62 to 0.9V vs Ag/AgCl^[25]^ and −0.6 to 0.8V vs Ag/AgCl^[17]^. These values are a function of the electrode material and the solution within which testing is conducted. For assessment in the context of new materials for neural stimulation, understanding the limits of the water window in physiologically relevant solution is important. Therefore, we performed a cyclic voltammetry sweep for silver wire and the Injectrode samples with an expanded range from −1.9 to 3V vs Ag/AgCl. It is important to note that, silver particles were used as the conductive filler for the Injectrode due to silver’s high conductivity and low cost for iterating through different Injectrode fabrication and testing methods to establish this early proof of concept. Despite these benefits, silver is not an acceptable choice for the creation of a chronic neural interface as silver and silver chloride have well understood toxic effects.^[26]^ Although testing performed with silver electrodes was sufficient for validating the Injectrode concept, future development will focus on the use of stainless steel, gold, or platinum filler particles that have been well validated in chronic neural interfaces.^[25, 27]^ We compared the Injectrode samples to a silver wire with the same geometric surface area to assess equivalency of electrochemical behavior over the expanded voltammogram but did not conduct a detailed study of the electrochemical behavior over this range.

As has been described elsewhere, in PBS, pure silver wire exhibited a large oxidation peak near 0V that has been attributed to the formation of silver chloride (AgCl)^[28]^, while the broadness of this peak and the observation of multiple humps may indicate the formation of other silver oxides.^[29–31]^ Additionally, the reversal of the silver chloride reaction was observed on the anodic side of the CV sweep (see Figure 5A). Putative hydrogen evolution was observed near the anodic limit of the voltammogram (at approximately −1.7V) and visually confirmed through the formation of microbubbles at the surface of the exposed silver wires and Injectrode samples. Interestingly, no clear oxygen evolution was observed even with the upper limit of the voltammogram sweep extended to 3.0V. This phenomenon was not explored further. One of the primary purposes of the electrochemical characterization experiments was to show that the Injectrode behaves similarly to the base metal, in this case silver. This was clearly true via visual inspection, although the Injectrode exhibited slightly decreased peak heights compared to a silver wire electrode with the same geometric surface area, peak shapes and locations for the Injectrode and base metal were similar in shape and location. This was the case for both the truncated sweep within the water window for platinum (Figure 5B) and the expanded sweeps performed on the silver wire and Injectrode (Figure 5A).

**Figure 5:**
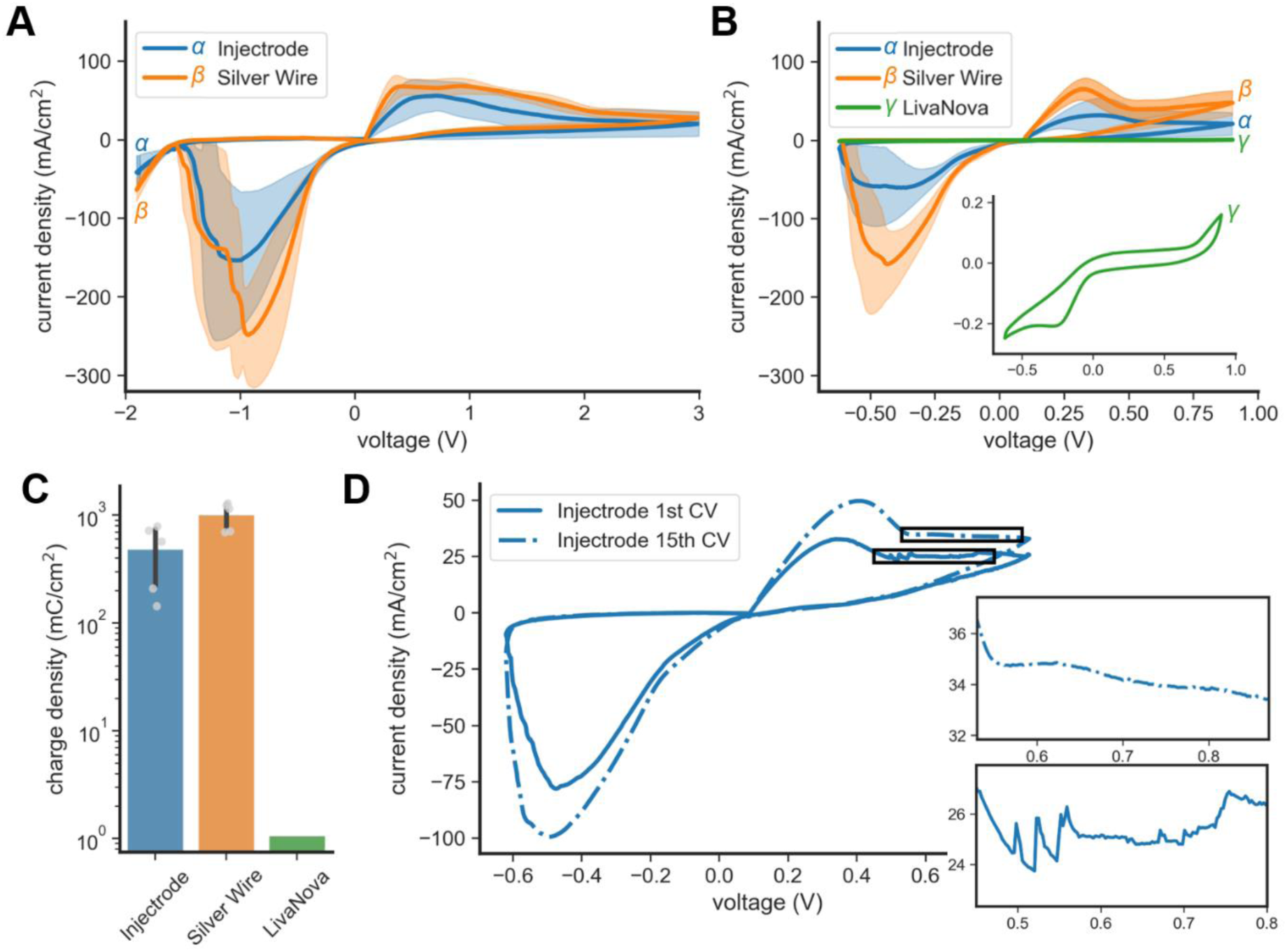
Cyclic voltammetry comparison. **A.** Mean cyclic voltammograms of five samples each of Injectrode and silver wire electrodes with matched geometric surface area. Scans were performed at 50 mV/s and over a range from −1.9V to 3.0V. **B.** Mean cyclic voltammograms of five samples each of Injectrode and silver wire electrodes as well as one contact of a LivaNova PerenniaFLEX^®^ lead. Voltammograms were scanned from −0.62 to 0.9V at a rate of 50mV/s. Inset show an expanded view of the voltammogram for the LivaNova electrode. **C.** Cathodic charge storage capacity (CSC_C_) calculated from the −0.62 to 0.9 V voltammograms for each set of electrodes. LivaNova and silver wire electrodes exhibit similar CSC_C_. **D.** The 1st and 15th voltammogram from one representative Injectrode sample showing spiking behavior in early voltammetry sweeps that goes away with the continued application of cyclic voltammograms. Insets show a zoomed view of this phenomenon. Shaded regions in panels A and B represent ± 1 standard deviation. Error bars in panel C also represent standard deviation.

Despite the expanded water window of Ag, a −0.62 to 0.9V vs Ag/AgCl voltammogram normally used for assessment of platinum was used instead of the expanded sweep for comparing the Injectrode voltammogram and charge storage capacity to that of platinum stimulation electrodes to provide a more fair comparison. The Injectrode and silver wire electrode both exhibited much larger CSC_C_ compared to a platinum electrode. This is largely due to increased charge injection through faradaic mechanisms. However, in light of the decrease in exposed silver available for faradaic reactions when looking at a two-dimensional cross-section of the Injectrode, as can be seen in the SEM imaging shown in Figure 4C and D, the similarity between the CSC_C_ of the silver wire and Injectrode samples was unexpected. This suggests that the Injectrode material has greater available three-dimensional surface area due to the previously discussed porosity and permeability of the matrix to the electrolyte to achieve greater CSC_C_. This effect is similar to the increases in CSC_C_ that have been observed with electrode coatings such as sputtering iridium oxide films (SIROF) and activated iridium oxide films AIROF on platinum.^[17,32–35]^

An interesting epiphenomenon was observed that was inconsistent between silver wire electrodes and the Injectrode samples. In extreme voltages of both the anodic and cathodic sweeps of the voltammogram, sharp peaks were initially observed during early CV sweeps (Figure 5D). These peaks are shown in detail in the lower inset of Figure 5D and are inconsistent with traditional faradaic reaction mechanisms that lead to ‘humps’ in the CV, such as those described for silver/silver chloride. Additionally, these spikes appeared at inconsistent voltages on subsequent CVs, and as additional CVs were run on the Injectrode samples the frequency of these spikes decreased. An example of this is shown in Figure 5D, where extensive spikes are shown on the 1st complete CV scan but are not observed on the 15th scan. These observations suggest the spikes are not caused by reactions occurring at specific voltages but instead may indicate changes in the exposed surface of the Injectrode or the presence of contaminants from the manufacturing process. We speculate that these spikes may be indicative of dissolution of base silver particles less well-bound within the silicone matrix, or of the exposure of new metal particles available for faradaic reactions through the opening of new pores in the polymer matrix. However, as the overall area under the CV tended to increase over initial sweeps and then settle into a consistent value, this would suggest the loss of particles or changes in exposed surface area would not negatively impact electrical characteristics of the Injectrode, from a neuromodulation perspective. Loss of metal particles from the surface of the Injectrode could be minimized, at least to some extent, through future refinement of the particle sizes, polymer matrix, mixing procedure, or other factors. However, it is worth noting that large chunks of platinum iridium have been reported in post-mortem human tissue from chronically stimulated cochlear implants and that tissue damage is driven by the formation of specific Pt salts during stimulation that lead to local toxicity^[24]^, as opposed to larger chunks of unreacted Pt or Pt/Ir flaking off.^[18,36–39]^ Whether loss of a small portion of conductive particles during the application of stimulation is an issue for strategies that leverage conductive particles to dope traditionally insulative matrices, such as the Injectrode, should be explored in future studies.

### 3.3. Electrochemical Characterization: Voltage Excursions

Although slow cyclic voltammetry provides unique information about electrochemical reactions that occur at the electrode/electrolyte interface, neuromodulation therapies typically employ charge balanced square wave current controlled pulses ranging from 30 µs to 1000 microseconds in duration. For that reason, voltage transients (or voltage excursions) are traditionally used to measure the polarization on the electrode itself during the application of cathodic leading, constant current, charged balanced biphasic pulses at a pulse width within normal clinical neuromodulation ranges^1^. For Pt/Ir electrodes, the current applied is increased until the cathodic polarization on the electrode itself approaches the cathodic limit for the evolution of hydrogen on a Pt/Ir electrode, and this current is adjusted for the geometric area of the electrode to determine the safe charge-injection limit. It is worth noting that for materials that differ from Pt or Pt/Ir the use of voltage excursions to determine unsafe levels of polarization needs to be reevaluated on an electrode material by electrode material basis. This is of particularly importance for materials through which there may be reversible safe electrochemical reactions induced at one level of polarization, and another unsafe electrochemical reaction occurring at a higher level of polarization.

Given the extended CVs performed to determine the water window for silver indicated the anodic and cathodic limits before the evolution of oxygen or hydrogen were outside the ranges of traditional Pt electrodes, and the use of silver particles within the composite material of the Injectrode is known to be an issue for direct translation, we decided to perform a conservative comparison of acceptable charge injection limits. This was done by intentionally limiting the acceptable polarization on the pure Ag wire or Injectrode to the established cathodic limit for Pt or Pt/Ir electrodes. By limiting the polarization of the electrode to the safe limits for Pt or Pt/Ir before hydrogen evolution we did not find the maximum charge injection limits of the Injectrode sample; however, this approach is sufficient to show the Injectrode concept can achieve neuromodulation relevant charge injection limits and is not inherently limited by the properties of the composite material. Given that the Injectrode silicone matrix is permeable to water (or PBS, or electrolyte solution *in vivo*) over time, the increased access to interior fractal dimensions during slow scan voltammetry may lead to a large CSC_C_ calculation that is not relevant for standard neuromodulation pulses. As such, voltage transient measurements provide an additional method to assess the electrochemical behavior during more stereotypical short neuromodulation pulses. For this reason, it will be important to determine safe charge injection limits for a more clinically accepted conductor/carrier composition in time.

As shown in Figure 6A, the voltage transient on the Pt/Ir Clinical LivaNova lead first consists of a very sharp negative voltage deflection at the initiation of the cathodic current pulse, resulting from the voltage generated from application of the current to the resistive load from the ionic solution and internal properties of the Injectrode. There is then a slow, consistent change in voltage during the remainder of the cathodic current pulse, generated by the polarization of primarily capacitive Pt/Ir electrode. Charge density limits before reaching −0.6 Volts of electrode polarization were between 50 and 150 µC/cm^2, consistent with the known charge density limits for Pt electrodes (Figure 6C). In comparison to the LivaNova leads, there is almost no polarization of the pure silver wire electrodes or the Injectrodes during the application of a current pulse adjusted to match the same applied current density (See Figure 6A-B). Overall, Injectrodes and silver wires behaved similarly during the applications of voltage transients, which is another indicator that the Injectrode will behave similarly to metal electrodes made from the same material as the conductive filler in the electrode. Small differences, between electrodes, in the initial voltage caused by the resistance of the solution are expected, as the two electrode types have slightly different geometric surface areas and geometries. The uncompensated resistance is driven primarily by the ionic resistance at the cross-sectional area of the electrode/electrolyte interface; since ionic current flow further from the electrode is not limited to the cross-sectional area of the electrode/electrolyte interface and take can advantage of the much greater volume for ionic current flow in the bulk solution. The internal resistance of the Injectrode, compared to the silver wire and LivaNova electrode may also contribute to these differences.

**Figure 6:**
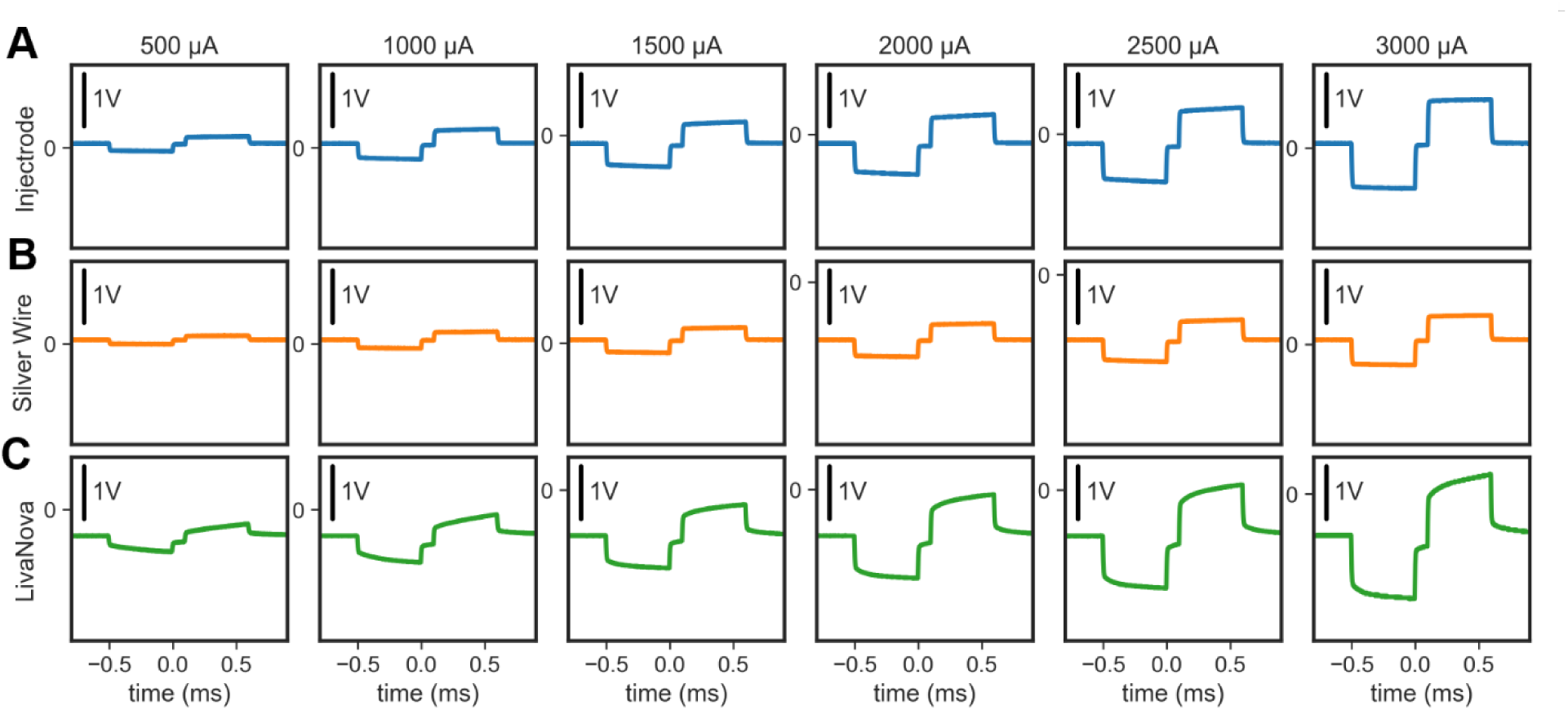
Voltage Transients. Voltage transients were performed in phosphate buffered saline using a 500µs pulse width and allowing at least 1000 pulses for the response to stabilize. A. Voltage transients performed at 500 through 3000 uA on one Injectrode sample that had previously undergone 20 cyclic voltammogram sweeps from −0.62 to 0.9V. B. The sample measurements shown in panel A but performed with a silver wire working electrode with geometric surface area matched to the Injectrode sample. C. Voltage transients performed with a LivaNova PerenniaFLEX^®^ stimulation electrode. To optimize visual comparisons of the progression of the voltage transients with respect to stimulation amplitude, the baseline for all voltage transients are centered on the vertical axis; however 0V with respect to Ag/AgCl is indicated on each vertical axis for assessing offsets.

### 3.4. Electrochemical Characterization: Impedance Spectroscopy

In addition to providing a relative measure of impedance at different frequencies for different electrode materials, electrochemical impedance spectroscopy (EIS) provides additional information about the balance of resistive vs capacitive effects dominating current flow. Figure 7A is an EIS sweep between 0.1 Hz and 100 kHz for the LivaNova lead. The Pt electrode EIS results can be explained using a simplified model assuming the Pt electrode/electrolyte interfaces acts as resistor and capacitor in parallel. This interface in turn is in series with the ionic resistance of the solution, and the impedance of the large Pt foil counter electrode is neglected due to the use of a three electrode setup. At very high frequencies in the 10-50kHz range, the capacitive element of the electrode/electrolyte interface is very low impedance with respect to both the resistive element of the electrode/electrolyte interface and the ionic resistance of the PBS solution. Consequently, the ionic resistance of the solution dominates the overall measured impedance as it is in series with the electrode, and the phase angle measured is 0 degrees indicating of current flow across a resistor. As the applied frequency decreases until another transition point 1-10 Hz, the impedance of the capacitive component of the electrode/electrolyte interface increases, and the phase angle increases towards purely capacitive current flow (90 degrees). As the applied frequency goes below 1 Hz the resistance component of the electrode/electrolyte interface becomes dominant with respect to both the impedance of the parallel capacitive component and the solution resistance, and the overall measured impedance begins to dramatically increase with the phase angle trending back towards purely resistive values (0 degrees).

**Figure 7:**
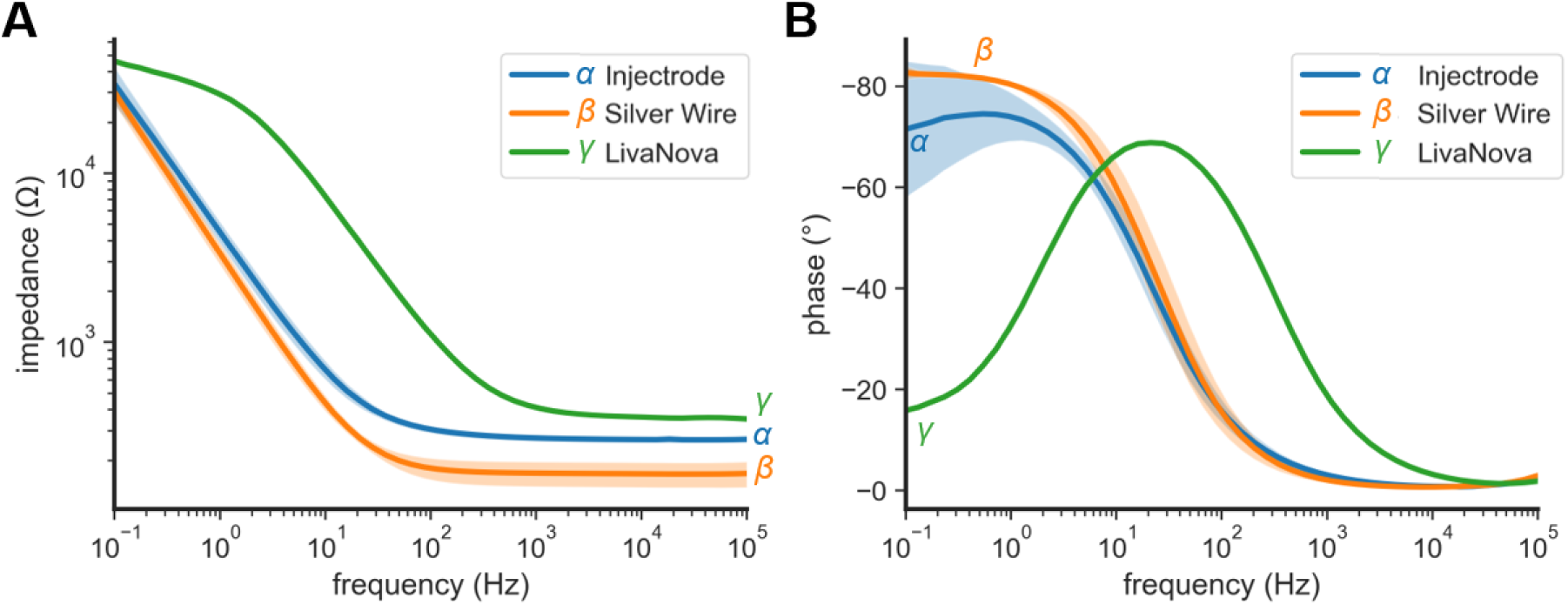
Electrochemical impedance spectroscopy comparison. Electrochemical impedance spectroscopy of five samples each of Injectrode and silver wire electrodes as well as one contacts of a LivaNova PerenniaFLEX^®^ lead. A. Mean impedance of each set of electrodes as a function of frequency. B. Mean phase of each set of electrodes as a function of frequency. Impedance and phase of Injectrode samples and silver wire show similar behavior across frequencies. Shaded regions represent ± 1 standard deviation.

For a pure silver wire the overall impedance at low frequencies is less than the Pt electrode, and the phase angle stays much closer to zero degrees above 100 Hz indicating current flow through a resistor (Figure 7B). This result is not surprising, as the silver wire has a much more facile mechanism for faradaic (resistive) charge transfer through the formation of AgCl than non-faradaic charge transfer (capacitive). The EIS curves for the Injectrode are very similar to those of a pure Ag electrode. The overall Injectrode impedance is slightly higher than the pure Ag wire at high frequencies, and the impedance becomes almost identical to Ag wire at low frequencies. This may suggest that more of the fractal dimensions of the silver particles are accessible to ions in the electrolyte solution through the silicone matrix at low frequencies. This low frequency behavior is a known phenomenon for conductive polymer coating strategies intended to increase the effective fractal dimensions of a stimulating electrode, decreasing pore resistance by providing more time for the ions in solution to permeate the inner recesses of the conductive polymer matrix during each sinusoidal pulse.^[32–34]^ However, given the complex nature of the Injectrode/electrolyte interface caution should be observed in overinterpreting these results.

### 3.5. Mechanical Testing

The Injectrode was capable of undergoing large reversible deformations (Figure 8). Of four samples tested, one initial sample was tested to only 140% strain (gray curve), the other three were tested to 200%, of which only one failed, at 167% strain. In comparison to the relevant neuroanatomy, the ulnar nerve is thought to undergo one of the largest deformations of nerves in the body and is only estimated to stretch by ∼29% strain under normal elbow flexion.^[40]^ The Young’s Modulus of the Injectrode was estimated from the slope of the elastic region in the stress-strain curves to be 72.1 +/- 10.9 MPa. This is orders of magnitude less stiff than other materials currently utilized in neural interfaces, e.g., steel (∼200 GPa), gold (74 GPa), polyimide (2.5 GPa), and PTFE (400 MPa).^[41]^

**Figure 8:**
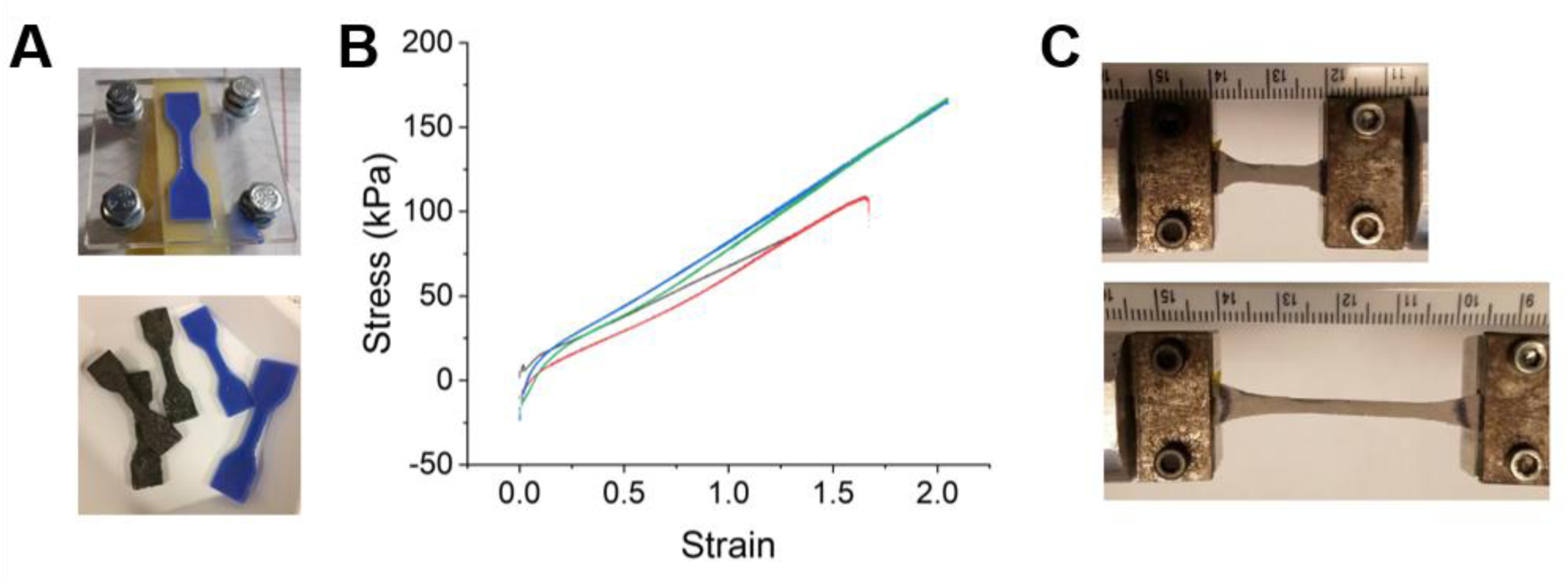
Tensile mechanical testing. A. Dog bone shaped samples of Injectrode material were prepared in acrylic molds (blue, Injectrode base material, gray, conductive silver Injectrode), modeled after ASTM D412-16, standardized testing of vulcanized rubber and other elastomeric materials. B. The elastic region of an Injectrode under tensile testing shown by a representative stress-strain curve. C. Photographs of Injectrode sample during testing, showing that the material is capable of elastically stretching to more than double its original length before plastic deformation occurred.

### 3.6. Surgical Delivery Proof of Concept

For proof of concept and visualization purposes, we demonstrated the ability to place Injectrodes with an open-cut down approach in a swine cadaver. The materials were easily extruded from 18 gauge needles affixed to syringes and formed cohesive and flexible structures around the nerves (Figure 9A). Importantly, the materials were capable of flowing around complex branched structures and conformed to the anatomy. The materials could also be injected into enclosed areas such as the tissue ‘sheath’ overlying the neurovascular bundle (Figure 9B). While these procedures were performed with an open incision, the modality could be adapted to a minimally invasive approach mimicking similar endoscopic procedures performed today.

**Figure 9:**
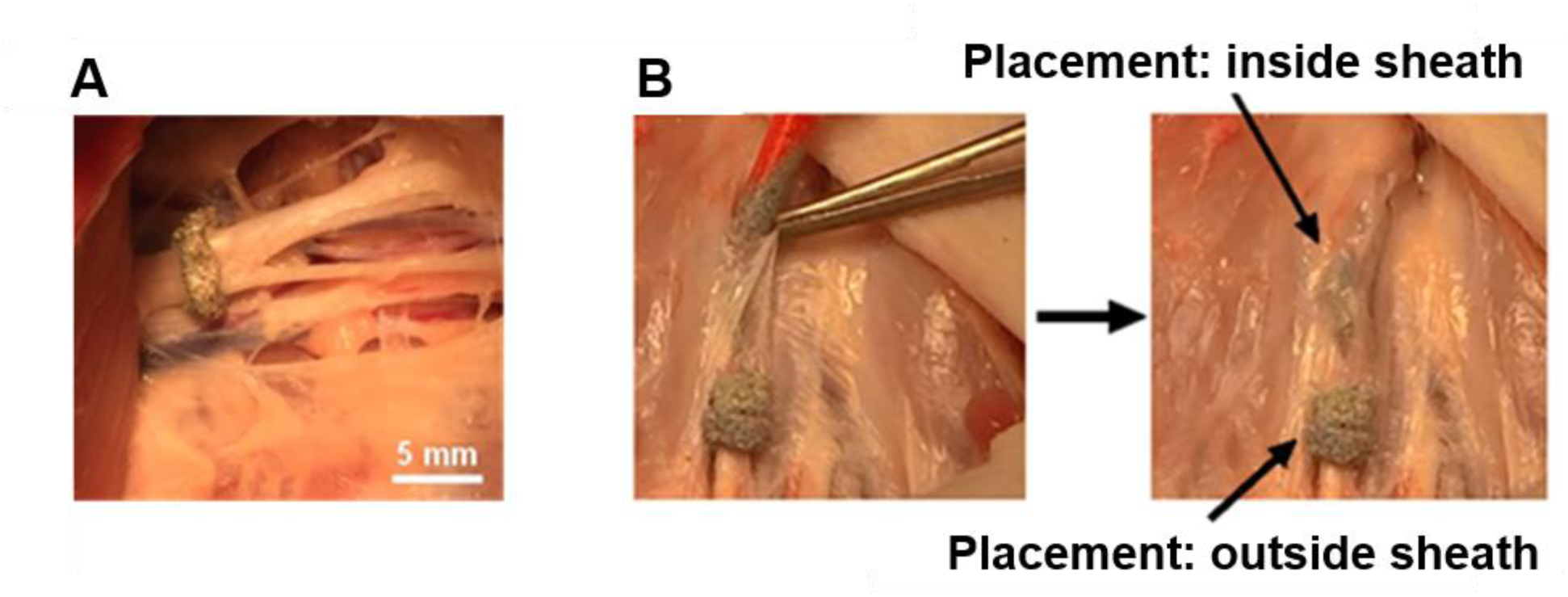
Delivery to Multiple Complex Nerve Anatomies in Swine. Injectrodes were used to stimulate nerves in a rat (N=5) and a swine (N=2) model following extensive cadaver testing to optimize delivery and easy handling under surgical-OR conditions. A. Delivered by syringe, nerve cuff diameters of 1 to 3 mm were achieved around peripheral nerves in a swine cadaver. In addition to encasing a single nerve with an injectable cuff, several nerves running in parallel - either in a bundle or at a neural plexus - were encapsulated and stimulated with a wire that was embedded within the Injectrode cuff. B. Furthermore, nerve bundles were encased by injecting the Injectrode into a nerve sheath commonly surrounding the nerve(s) and then gently manipulating (massaging) the nerve sheath, thereby distributing the Injectrode inside the sheath that functioned as a mold. Injectrodes were placed around nerves on the inside and on the outside of the nerve sheath in swine model.

### 3.7. Electrical Stimulation of Compound Motor Nerves (Brachial Plexus)

Injectrode cuffs were placed 360-degrees around the brachial plexus in a rat model, just distal to the location where the nerve visibly branched in to the median, ulnar, and radial segments. A silver wire was embedded into the Injectrode cuff prior to curing and used to connect to the signal generator. A nerve recruitment curve was collected and supramaximal recruitment was determined to be beyond 2 mA of current amplitude. To ensure a stable system during tetanic stimulation experiments, at least twice the supramaximal current amplitude was used, in this case 5 mA pulses of 200 µs pulse width. Complete tetanic muscle contractions resulted from 30 Hz stimulation using a charge balanced biphasic waveform (200 µs pulse width, 5 mA supramaximal amplitude) as shown in Figure 10. The simulation effect was quantified by direct measurement of joint angles in captured video (Figure 10A).

**Figure 10:**
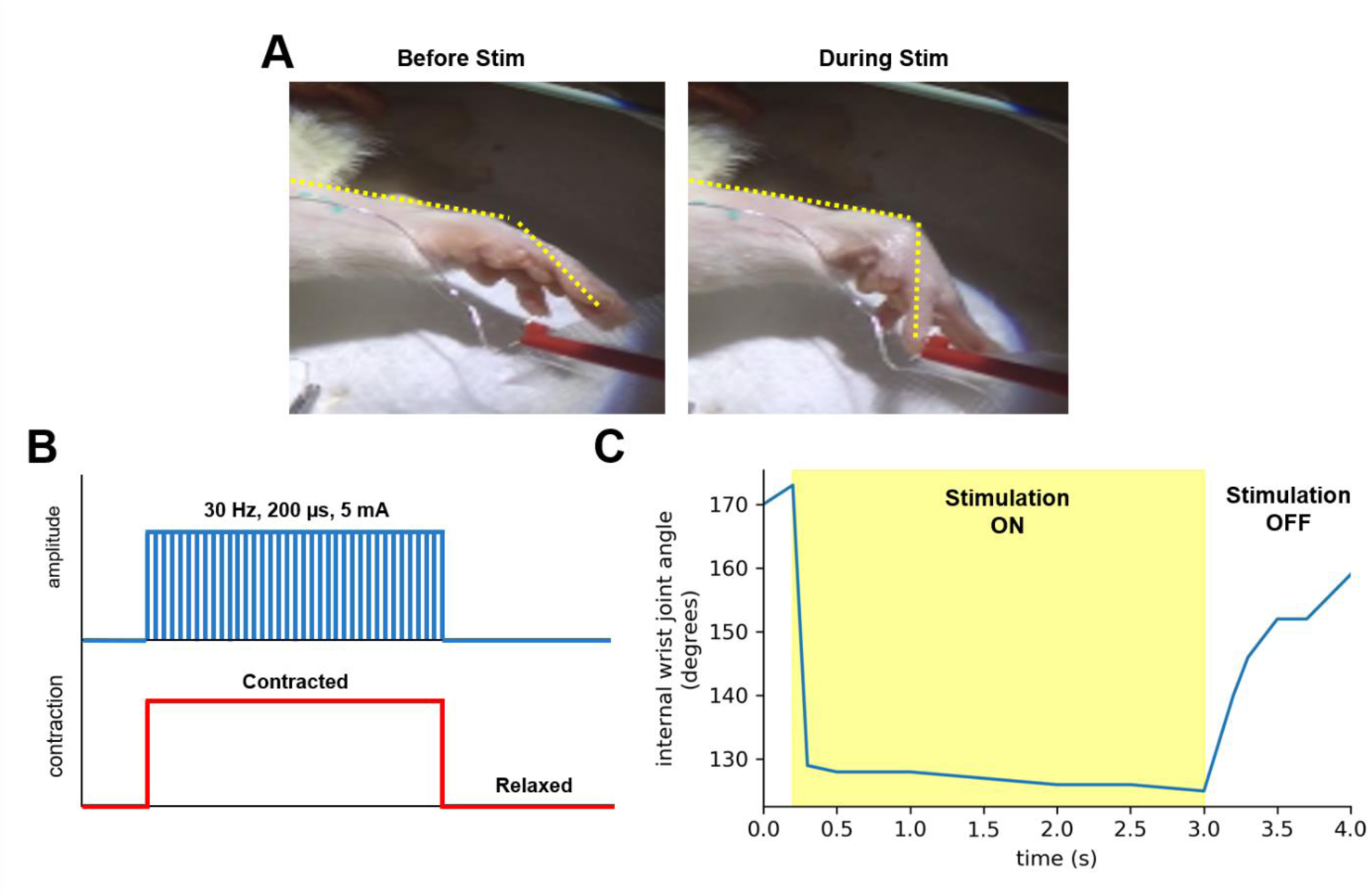
Acute Tetanic Stimulation Rat Brachial Plexus. Injectrodes were placed in a bipolar arrangement on the left brachial plexus of a rat. A. Based on visual assessment, stimulation parameters were established to produce tetanic contraction of the forearm (30 Hz, 200 us, 5mA). B. A period of stimulation was applied to produce a contraction. C. Contraction was and later quantified by measuring wrist joint angles via post-hoc by video analysis.

This brief experiment demonstrated that the Injectrode material can form an electrically conductive interface with the nerve and produce activation of the fibers that can lead to a tetanic contraction of muscles. In addition, we demonstrated that the material could form an interface with the complex branched nerve structures of the brachial plexus.

### 3.8. Electrical Stimulation of Swine Vagus Nerve

In order to directly compare stimulation thresholds with as closely controlled electrode geometries as possible, we formed the Injectrode materials into bipolar cuffs of the same size and electrode spacing as the clinically available LivaNova cuff electrode (Figure 11A). We performed a current controlled dose-titration protocol, where the biphasic waveform was held constant, 200 µs pulse width at 30Hz stimulation, and current was ramped up between 0 to 5 mA. The physiological parameter we measured was heart rate (BPM), per previously published protocols.^[42]^ For a given amplitude, the Injectrode induced a proportionally lower change in heart rate compared to the LivaNova cuff electrode (Figure 11B). The approximate voltage was estimated based on recorded cuff impedance values prior to stimulation. Interestingly, though not surprisingly, when corrected for voltage titration, the two electrodes performed nearly identically, as shown in Figure 11. which were placed in the same location and configuration on the vagus nerve. B. Higher stimulation currents were required with the Injectrode compared to the LivaNova to drive the same heart rate response. C. However, Impedance of each electrode was measured to be 1350-ohm for the LivaNova and 360-ohm for the Injectrode. Correcting for lower impedance of Injectrode connection, dose-titration on Voltage-BPM scale was nearly identical with one another, demonstrating a characteristic sigmoidal response relationship. In panels B and C spline curves were fit to each dataset to improve visualization of the relationship between stimulation and response.

**Figure 11:**
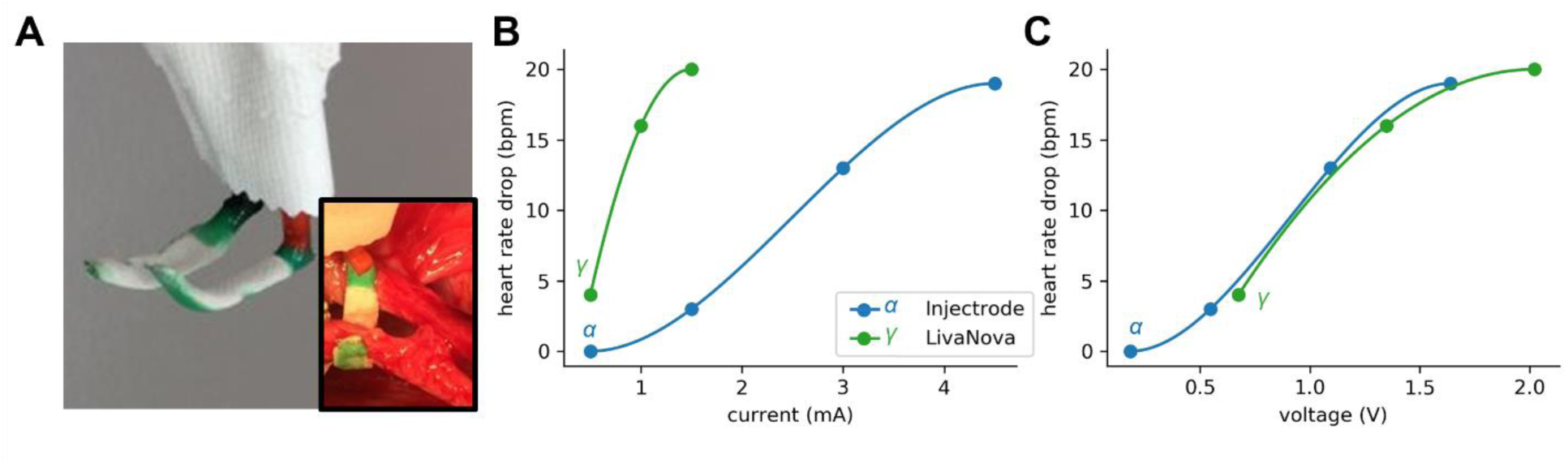
Demonstration of Acute Swine Vagal Nerve Stimulation. To evaluate effective stimulation parameters of the Injectrode compared to the commercial LivaNova lead to induce heart rate reduction, Injectrodes were placed in an open-cut down procedure on the vagus nerve. To limit variables of the comparison, Injectrodes were fabricated outside the body A. Stimulation dose-titration was performed with a current-controlled simulator. The experiment was performed first with a standard LivaNova cuff, followed by bipolar Injectrode contacts

### 3.9. Benefits, Challenges and Future Directions of the Injectable Electrode

The Injectrode is a unique electrode technology that can be injected onto, into or around a target structure, where it cures in vivo to form a conductive tissue interface. The fluidity of the pre-polymer material when it is injected allows for the electrode to conform to the patient’s individual neuroanatomy and thus provides a scalable approach to the size and geometry of different target structures. This gives the Injectrode the potential to simplify the surgical approach and eliminate complicated electrode designs needed for difficult neuroanatomical targets such as the stimulation of a nerve plexus presented in Figure 10. Additionally, the elastomeric properties of the cured material has a Young’s modulus of under 100 kPa. While this is substantially stiffer than the neural tissues, it is still orders of magnitude less stiff than traditional neural interface materials made from solid wires with polymer insulation. A reduction in the mechanical mismatch between the device and neural tissue may reduce cyclic strains and stresses on the device and surrounding anatomy, and consequently improve the reliability of a chronic neural interface.^[43, 44]^ Despite the benefits derived from the material properties of the Injectrode there are a number of design considerations that will need to be addressed by future studies, including the use of conductive metal particles with known chronic biocompatibility such as gold, platinum or stainless steel.

A number or previous studies have taken advantage of silicone (PDMS) composites doped with conductive filler particles in order to create a conductive material^[45–47]^, including for use the developement of neural electrodes^[48]^. Additionally, other in-body curing PDMS composite materials have been proposed in the literature^[49, 50]^ and are commonly used in small animal studies^[51–54]^ due to PDMS being physiological inert and favorable curing properties; PDMS cures in an aqeous environment without the formation of toxic products or exothermic heat^[49]^. These advancements in conductive and in-body curing polymers have not been applied to create an injectable electrode, but they provide a basis for the use of silver particles as a conductive filler within a stretchable silicone substrate. Based on these previously published studies, and the low cost of silver for iterating through the proof of concept experiments presented in this paper, in the present proof of concept study silver particles were selected as the conductive filler in the composite material. Despite its use as an antimicrobial in certain medical applications, in high doses and during long-term contact with tissue, dissolution of silver metals and the formation of Ag salts, is well known to be associated with cellular and organ toxicity.^[26]^ As such, the exact Injectrode formulation used in this paper is not viable as a chronic neural interface. To overcome this limitation, we are currently testing Injectrode formulations based on the use stainless steel, gold, platinum and other non-metallic filler particles to provide conductivity. Due to the need for future changes in the conductive filler used in the composite Injectrode material, we focused the electrochemical testing presented in this paper on demonstrating similarity to the base conductor (silver) and showing that the composite material construct is capable of neuromodulation relevant charge storage capacities and charge injection limits. This was sufficient for this proof of concept demonstration of feasibility, in future studies we intend to emphasize developing the neural interface in terms of both bio-electrochemical compatibility and material flexibility.^[55]^

Another important point of future development is the Injectrode implantation procedure. While the experiments presented herein were performed through an open cut-down to the neuroanatomical target, the ultimate goal of the Injectrode concept is to enable the completion of these procedures through a minimally invasively surgical approach. Achieving this goal will require refinement and further development of surgical delivery tools and image guidance tools for accurately injecting multi-contact Injectrodes in way that well-controlled and repeatable across surgeons. A particularly important aspect of non-invasive delivery will be the indirect visualization of the nerve target and associated instrumentation for accurately injecting the electrode followed imaging of the final Injectrode alignment for confirmation of target engagement. Typical imaging techniques for lidocaine nerve blocks utilize ultrasound imaging surrounding anatomical landmarks, e.g., bony structures and muscle fascia planes. Similarly, the implantation of spinal cord stimulation electrodes utilizes fluoroscopic guidance of the electrode. For this reason, intraoperative ultrasound and fluoroscopic methods for visualizing the procedure are currently being explored; other imaging modalities may also be useful for intraoperative guidance or postoperative verification of targeting. In addition to implantation of the conductive Injectrode, non-conductive polymer may also need to be delivered as insulation in order to isolate Injectrode contacts or direct current flow to the target of interest. However, the individual procedures for administration of conductive and non-conductive materials to isolate the neuroanatomy of interest will need to be optimized for specific indications.

The principal ingredients of the Injectrode were selected to provide rapid proof of concept, and because they have been used in different forms within other previously FDA-approved implantable products. Although the FDA does not approve specific materials, as performance depends on specific use in the context of an entire system, familiarity for implantable use can streamline the regulatory process. Similarly, surgical glues that cure within the body are not uncommon in clinical use and provide a model to follow for conductive Injectrode based solutions.

### 3.10. Advantages and Disadvantages of the Injectrode Compared to Other Recent Innovations in Soft Electrodes

Over the past decade, there has been a renewed interest in the use of thin-films and other ‘soft’ electrode materials to create high-channel, yet low geometric area electrodes with a Young’s Modulus closer to native tissue to minimize initial surgical trauma and the chronic immune response.^[54–60]^ These electrode development strategies include the use of PDMS substrates with conductive traces and electrode contacts to develop stretchable multi-contact electrode arrays for stimulation and recording.^[48,61–64]^ Additionally, shape memory polymer (SMP) or other ‘shuttles’ have been used to create a stiff carrier to facilitate implantation of electrodes into the brain, yet leave a soft electrode chronically to minimize the immune response of tissue.^[54, 58, 60]^ Similarly, the Charles Lieber group have demonstrated that a ultraflexible open mesh electrode array can be injected into the rodent brain via syringe and minimize the chronic immune response, enabling high quality chronic recordings for periods of at least 12 weeks.^[65]^ More recently, the Romero-Ortega group has demonstrated that SMP cuffs can be surgically implanted directly on the rat sciatic and pelvic nerves and both record and intermittently stimulate for periods of at least 30 days, while minimizing the immune response compared to stiffer control devices.^[66]^ In comparison to the Injectrode, these promising technologies have the clear advantage of higher channel counts, as well as predictable electrode geometries and spacing.

However, extensibility of these technologies to an embodiment compatible with minimally invasive delivery outside the brain yet capable of high-duty cycle continuous stimulation common for clinical neuromodulation therapies remains unclear. At present the only high-density thin-film electrode technology that is FDA market approved for over 30 day use is the Argus II Retinal Prostheses from Second Sight. Thin film polymers are permeable to water, and tend to fail chronically as fluid ingress finds pinhole defects between depositions layers which eventually lead to adhesion failures between layers and ultimately insulation between adjacent electrodes.^[67–70]^ Electrical stimulation through these devices exacerbates the failure modes by two mechanisms. First, electrical stimulation through an ultrathin platinum or platinum/iridium electrode causes transient mechanical deformation of the electrode, leading to lack of conformance to the insulation and failure of the lead traces connecting to the electrode.^[71–73]^ Second, the application of a DC bias for active electronics or normal charge-balanced constant current stimulation is known to hasten device failure, putatively by exacerbating fluid ingress through the polymer layers.^[67, 69]^ As of the writing of this manuscript, the long-term chronic viability of the Injectrode is also unproven; however, the Injectrode results presented here demonstrate that the metal particle-based strategy performs electrochemically under stimulation like a pure wire of the same material. This suggests that the chronic Injectrode behavior will resemble that of already clinically proven gold and platinum metal electrodes of similar geometric area.

By comparison to existing soft polymer electrode strategies, the Injectrode has the potential advantages of being very simple and inexpensive to manufacture. This is particularly important for viability in translation, as insurance payers determine the reimbursement price for implantable neuromodulation technology. Complexity increases the number of failure points, complicates the supply chain for manufacture, and ultimately increases the overall cost of manufacture. This is why simple implantable devices based off variants of the cardiac pacemaker but modified for the cervical vagus, deep brain, or spinal cord currently dominate the existing neuromodulation industry. Unlike the existing soft polymer strategies which have to be modified based on surgical approach and intended neural target, the Injectrode as a platform technology is extensible to injectable surgical deployment around any complex neural structure without modification to create highly conformal electrode interfaces. This also potentially enables unique uses such as injectable delivery of undoped Injectrode to insulate nerves implicated in therapy limiting off-target effects, or to ‘reconnect’ existing clinical leads that, through scarring or migration, no longer interface efficiently with their intended neural targets. Consequently, we envision both the Injectrode and high-density ‘soft’ thin film electrodes will likely each have unique uses and therefore their own ‘niche’ in the quickly evolving neurotechnology clinical market.

## 4. Conclusion

The Injectrode addresses problems with invasiveness, complexity, and cost of the implantation procedure that hinder the adoption of neuromodulation therapies and reduce patient’s access to these otherwise promising treatments. In order to develop the Injectrode into a chronically implanted neural interface, optimization of material properties and surgical approaches will need to be addressed in future studies. Additionally, long-term electrochemical testing of the Injectrode and histological analysis showing chronic stability and biocompatibility of the implant will need to be performed. However, the electrochemical testing and proof of concept experiments presented in this paper demonstrate the feasibility of the injectable electrode approach. Electrochemical testing, performed based on common FDA preclinical benchtop tests, including scanning electron microscopy, cyclic voltammetry, electrochemical impedance spectroscopy, and voltage transient analysis showed that the Injectrode, which is conductive due to the incorporation of metal filler particles into a composite material, is electrochemically similar to metal wire of the same material (silver). Additionally, acute *in* vivo testing of an Injectrode, cured around the complex compound motor nerve branches of the brachial plexus, demonstrated stimulation induced tetanic activation of the terminal muscles. Finally, in an animal model better matching the scale of human anatomy, proof of concept comparisons of Injectrode performance to a clinical LivaNova vagus nerve stimulation lead showed stimulation induced heart rate changes in the swine. These experiments suggest the feasibility of the Injectrode for neural stimulation. Additionally, by virtue of being simpler than traditional electrode designs, less invasive, and more cost-effective, the Injectrode has the potential to increase the adoption of neuromodulation therapies and enable the treatment of new indications.

## Acknowledgements

The authors would like to acknowledge funding from The Defense Advanced Research Projects Agency (DARPA) Biological Technologies Office (BTO) Targeted Neuroplasticity Training Program under the auspices of Doug Weber and Tristan McClure-Begley through the Space and Naval Warfare Systems Command (SPAWAR) Systems Center with (SSC) Pacific grants no. N66001-17-2-4010.,

We would also like to acknowledge Erika Woodrum and the FES Center for their help in constructing Figures 1 and 2, and to Janet Gbur and John Lewandowski at the Advanced Manufacturing and Mechanical Reliability Characterization Center for their assistance in collecting mechanical testing data.

Additionally, we gratefully acknowledge use of facilities and instrumentation supported by NSF through the University of Wisconsin Materials Research Science and Engineering Center (DMR-1720415).

Lastly, we would like to thank Seth Hara from the Mayo Clinic for his feedback on this manuscript.

## Footnotes

1 Cathodic leading pulses are traditionally used for voltage excursion tests, as cathodic leading biphasic pulses require lower thresholds for activation when an electrode is placed directly on a nerve fiber. When there is any separation between the electrode and the fibers, or a group of fibers are being stimulated that cover a span of distances from the electrode, there may be less of an advantage to the use of cathodic first pulsing strategies.

