## Supplementary Information for "A Truly Injectable Neural Stimulation Electrode Made from an In-Body Curing Polymer/Metal Composite"

### Methods - Bulk Conductivity (Dry Bench Testing)

Electrochemical testing was performed to determine material characteristics applicable to the use of the Injectrode for neural stimulation. Conductivity of bulk material was determined for a range of silver/silicone ratios. Preliminary blends of polymer and silver were generated and optimized for ability to easily flow through a standard 1 cc syringe, low impedance at 1 kHz, and ability to fully cure in under 5 minutes within thawed chicken. The preliminary testing was used to inform the experimental design sample compositions.

In order to test the conductivity of the electrode using samples with controlled material properties and geometry, components of the Injectrode composite were combined in a 1 cc syringe and mixed for 30 seconds to make the flowable pre-polymer. To control the geometry, the pre-polymer was immediately injected into individual wells of a 96-well plate with coiled silver wire contacts placed at the top and bottom of each Injectrode sample. After allowing 10 minutes for the sample to cure, impedance of each sample was measured using 1 kHz sinusoid from a standard LCR meter (DE-5000, DER EE Electrical Instrument).

### Results - Bulk Conductivity (Dry Bench Testing)

Initial bench testing of electrode impedance at 1 kHz was performed on dry samples using a LCR meter (Figure4 A,B), and demonstrated that the composite material is low impedance (<10 Ω), above a percolation threshold of approximately 65% w/v mix with respect to silver content.

In contrast, samples with silver content below the percolation threshold maintained a high impedance greater than 10 MΩ (Figure 4C). Injectrodes are electrically conductive immediately following placement; curing and cure times of 90-120 seconds were observed with the formulations tested. However, the impedance measurements presented here were all performed after allowing 10 minutes for the Injectrode samples to fully cure.

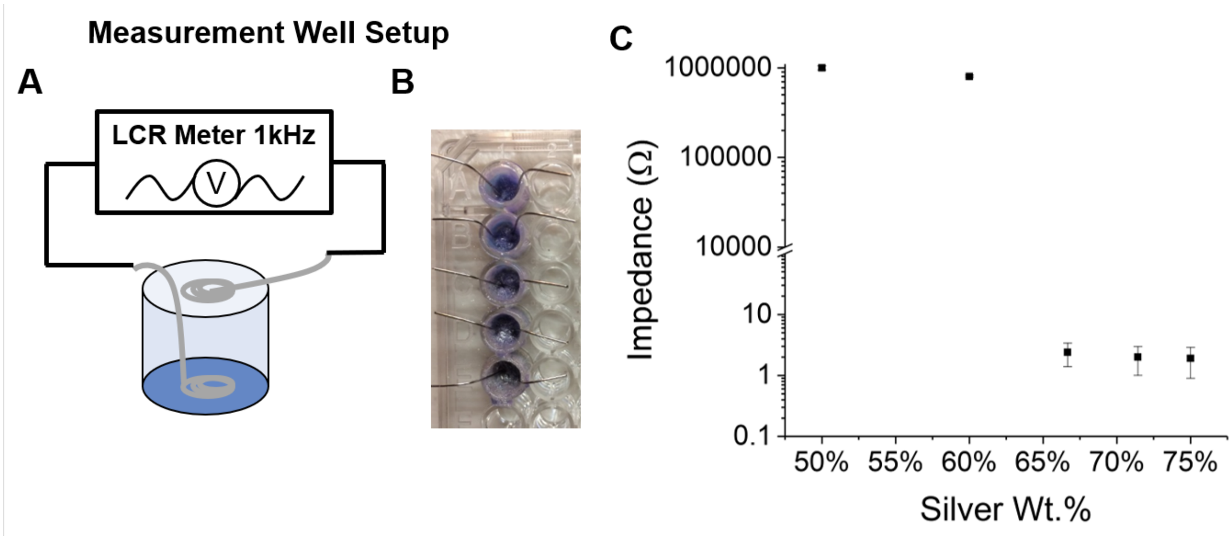


**Figure S1: Conductivity Versus Filler Weight Percent.** Composite Injectrode materials were prepared using various filler wt.% from 50-75% of total composition. **A.** After mixing, each material was extruded into a 96-well plate to control geometry and size. Before extrusion, a silver wire contact was placed in the bottom of each well to serve as one of the impedance meter contacts. A second silver wire contact was placed on top of the material in each well. After curing, impedance was measured using an LCR meter in ambient air conditions with a 1 kHz sinusoidal waveform. **B.** A photograph of the sample setup. **C.** A steep inflection point around 65 wt. % is suggestive of achieving percolation (a continuous network) of silver particles in the composite.

Percolation is a well-studied phenomenon that describes the material percentage required to forms an interconnected network throughout the composite, i.e. metal-to-metal network.[25,26] The percolation threshold is dependent on the volume of the particulate filler as well as the aspect ratio of the individual particles.[25,26] For simplicity of measurement during fabrication, weight ratios were the easiest to ascertain and control, but the use of weight ratios for controlling of impedance of the Injectrode is dependent on the use of consistently sized particles with similar distributions and dispersions. While these initial benchtop impedance measurements provided the ability to rapidly iterate through different formulations, they do not adequately represent the electrode performance under physiological use conditions, e.g. hydrated in a saline environment, during the application of electrical stimulation, etc. Therefore, electrochemical testing was better to assess the capabilities of the Injectrode for neural stimulation.
